# Automated Human Habenula Segmentation from T1-weighted Magnetic Resonance Images using V-Net

**DOI:** 10.1101/2022.01.25.477768

**Authors:** Joo-won Kim, Junqian Xu

## Abstract

The Habenula (Hb), a small bilateral midbrain structure, plays an important role in aversion and reward processing. Despite increasing interest in imaging human Hb structure and function, it is challenging to define the Hb in magnetic resonance imaging (MRI) due to its small size and low anatomical contrast to the surrounding thalamus. In this study, we developed a deep learning-based human Hb segmentation model. This model was trained and tested using about a thousand subjects’ 3T T1-weighted images with Hb labels from our previous myelin content-based Hb segmentation. The predicted Hb segmentation showed high similarity and small surface distance to the label; the average (across test subjects) dice similarity coefficient, mean distance, and Hausdorff distance were 0.79, 0.22 mm, and 1.63 mm, respectively. We also demonstrated out-of-sample robustness using other 7T and 3T MRI datasets. The proposed Hb segmentation method is automated, objective, fast, robust, and reliable, relieving the requirement of T2-weighted images in the established myelin content-based Hb segmentation to suite a broader range of MRI studies.

## 1. Introduction

The Habenula (Hb) is a small bilateral midbrain structure located next to the dorsomedial thalamus and the third ventricle (Namboodiri et al., 2016). The Hb consists of two subregions, medial and lateral Hb, which are functionally and anatomically distinct (Benarroch, 2015; Díaz et al., 2011). The Hb plays a crucial role in aversion and reward processing, and growing evidence suggests its implication in psychiatric disorders including depression and addiction (Fore et al., 2018; Hikosaka, 2010). This led to an increasing interest in human Hb structure and function research recently.

However, in vivo human Hb imaging is challenging because of its small size (about 30 mm^3^ in each hemisphere) (Ranft et al., 2010) and low anatomical magnetic resonance imaging (MRI) contrast to the surrounding thalamus in any single structural MRI modality, e.g., T1-weighted (T1w) or T2-weighted (T2w) images. However, its contrast to the thalamus can be increased by combining high-resolution 3D T1w and T2w images into T1w-over-T2w ratio (T1w/T2w) images due to its high myelin content (Glasser et al., 2013). Leveraging this enhanced Hb contrast in T1w/T2w image, we have developed an automated Hb segmentation method using this increased contrast, saying myelin content-based segmentation (Kim et al., 2018, 2016). This method can be applied to either functional (Ely et al., 2019, 2016) or volumetric(Jha et al., 2021; Schafer et al., 2018) analysis.

The requirement of high-resolution 3D T1w and T2w images of our T1w/T2w segmentation method, nevertheless, limits its wider applicability, because not until recently, the vast majority of neuroimaging research projects only acquired T1w anatomical images. Manual Hb segmentation (Carceller-Sindreu et al., 2015; Savitz et al., 2011; Schmidt et al., 2016) or geometric Hb segmentation using surrounding landmark structures (Lawson et al., 2013) have been widely employed if only T1w images were available. However, these manual Hb segmentation methods are laborious and subjective. Recently, other automated Hb segmentation methods utilizing only T1w images have been proposed, such as registration-based multiatlas-driven Hb segmentation (Germann et al., 2020) and deep learning-based Hb segmentation (Lim et al., 2021). Germann’s registration-based method depends on the quality of the registration to their atlases and takes a long process time. Lim’s deep learning method was trained and tested using their 69 participants’ axial 7T T1 map slices containing Hb, and therefore, it is more likely a project-specific model requiring 7T T1 maps and manual pre-selection of slices.

Deep learning has been widely applied to various image segmentation applications because of its versatility, remarkable performance, and short processing time. There has been much advancement in medical image segmentation using deep learning for small anatomical structures such as spinal cord gray matter 2D segmentation (Perone et al., 2018). U-Net is the most prevalent 2D image segmentation deep-learning model, and it has shown good performance on medical image segmentation (Ronneberger et al., 2015). However, segmenting small tear-shaped structures like Hb in 3D is much preferred to 2D. Application of 3D deep learning segmentation is still rare in medical imaging, probably due to a lack of reference segmentation for these small structures for training 3D deep learning models.

Deep learning-based segmentation requires, in general, more than thousands of training data to achieve high accuracy and to avoid overfitting because of its complex network structure. It is difficult to create several such labels (i.e., reference segmentation) from medical imaging data since many will require manual work. In order to solve this limitation, image data augmentation is commonly used by slightly modifying image data to increase the amount of data. Such augmentation may include flipping images, adding noises, transforming images, zooming, and so on. Although these methods artificially increase the number of labels, one still needs a large number of subject data to address subject-wise nature variability. By leveraging a large amount of readily available reference 3D Hb segmentations using our T1w/T2w segmentation method (Jha et al., 2021), we are uniquely positioned to use these for 3D deep learning model training.

Here, we introduce an automated, objective, fast, and robust Hb segmentation on T1w images using 3D deep learning to overcome the limitations of previous methods. We adopted V-Net (Milletari et al., 2016), a 3D version of U-Net, and trained the model with about a thousand reference Hb segmentation and evaluated its performance. In addition, we applied the trained model to other datasets to show its robustness.

## 2. Methods

### 2.1. Subjects and MRI acquisition

Training, validation, and test data were obtained from the Human Connectome Project Young Adult (HCP-YA) S1200 (Van Essen et al., 2012). The HCP-YA S1200 participants were healthy subjects (n = 1113) between 22-37 years old. 3T T1w and T2w structural images were acquired at 0.7 mm isotropic resolution (Glasser et al., 2013).

Other datasets to evaluate the Hb segmentation performance included CamCAN data (Shafto et al., 2014; Taylor et al., 2017), and Mount Sinai 7T data. The CamCAN data are of 652 subjects with a wide age range (18-88 years), and myelin content-based segmentation is applicable to them. They contain T1w and T2w images with 1 mm isotropic resolution from 3T Trio (Siemens) with the following parameters; T1w: 3D MPRAGE, TR/TE/TI = 2250/2.99/900 ms, FA = 9 °; T2w: 3D SPACE, TR/TE/TI = 2800/408/900 ms. The Mount Sinai 7T data contain MP2RAGE (Marques et al., 2010) UNI images with 0.5 mm isotropic resolution from 7T Magnetom (Siemens) with the following parameters; TR/TE/TI1/TI2 = 5000/5.75/900/2780 ms, FA1/FA2 = 5/3 °, covering most of the brain.

### 2.2. Reference Hb segmentation

To use as training, validation, and test labels for the deep learning process, we applied the previously proposed fully-automated version of myelin content-based Hb segmentation (Kim et al., 2018) from AC-PC aligned T1w, T2w, and T1w/T2w images. The resulting segmentations were probability maps ranging from 0 to 1. We segmented Hb from the HCP-YA S1200 and CamCAN data, and then we visually inspected the segmentation to exclude unsatisfactory segmentation such as underestimation or overestimation (Jha et al., 2021; Kim and Xu, 2018).

### 2.3. Machine learning setup

See Fig. 1 for an overview of our machine learning training and prediction process

**Figure 1.**
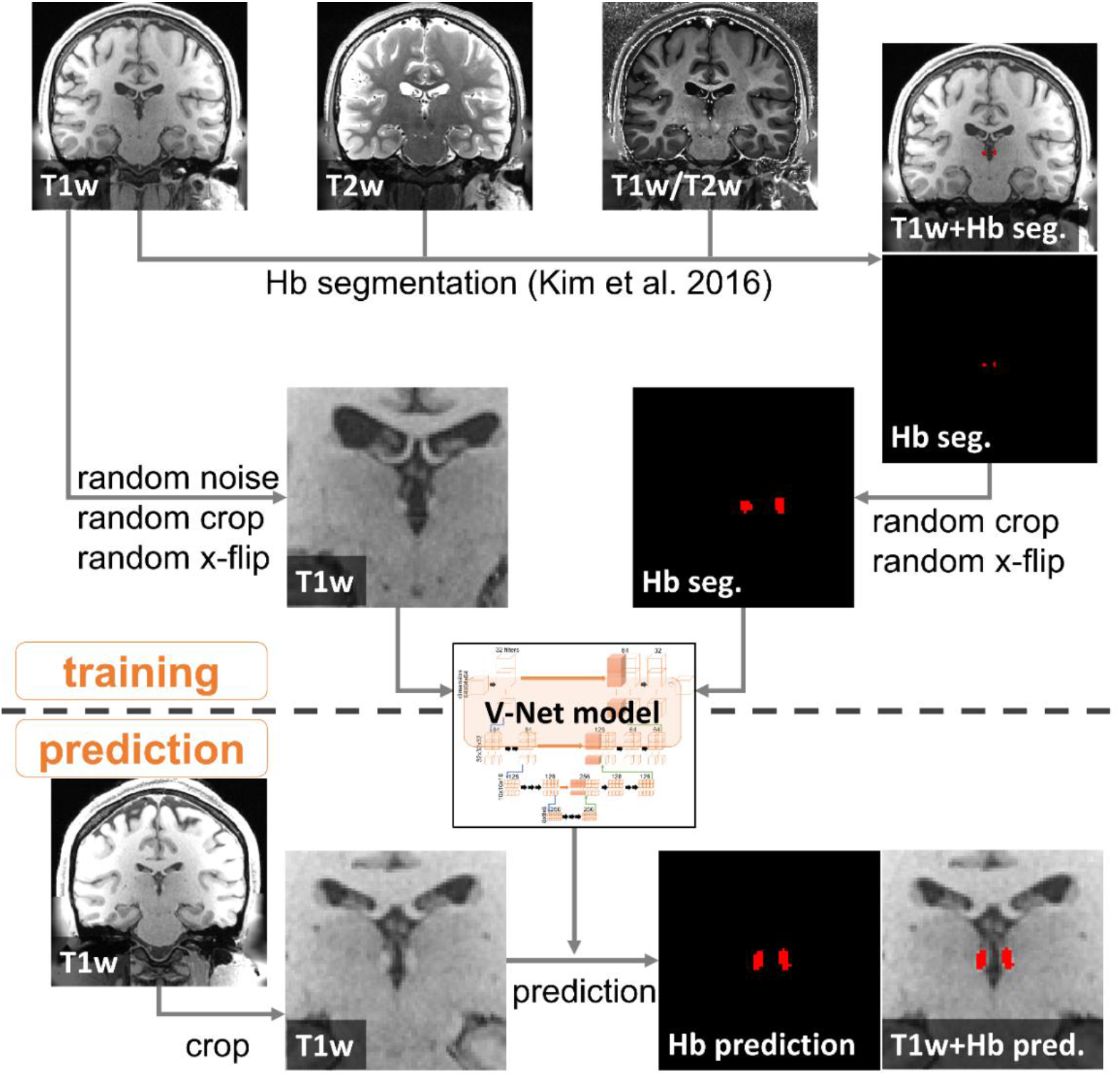
Overview of deep learning model training and prediction. Top: training process including Hb segmentation (Kim et al. 2016 using T1w, T2w, and T1w/T2w images) and image augmentation (random noise, crop, and x-flip). Bottom: prediction process including crop, prediction, and postprocessing.

We randomly separated the HCP-YA S1200 data with satisfactory Hb segmentation into training, validation, and test sets (n = 717, 127, and 157, respectively). The inputs were AC-PC aligned T1w images of the subjects, and the labels were the corresponding Hb segmentation after applying empirical threshold 0.3 to the Hb segmentation probability maps.

To reduce the input image size for memory limitation, we cropped the T1w images to 64×64×64 voxels centered at random locations around the Hb (center x, y, and z coordinates were within 110-146, 129-165, and 87-123 ranges, respectively, in the total 260×311×260 voxel space). To increase the number of training sets, we employed some image data augmentation methods during the training process. First, both with- and without-bias-field-correction (Glasser et al., 2013) T1w images were used which have the same shape but different intensities. Second, the center of the crop was random, resulting in the shift of the Hb location. Third, the images were randomly left-right-flipped. Fourth, random noise was added.

While a typical deep learning-based image classification network monotonically decreases the image dimension (matrix dimension), V-Net or U-net contains steps to recover the matrix dimension (Fig. 2, expansion) after decreasing steps (Fig. 2, contraction). At each recovering step, corresponding decreasing step’s matrices are concatenated. Through these recovering steps, the output can have the same dimension as the input.

**Figure 2.**
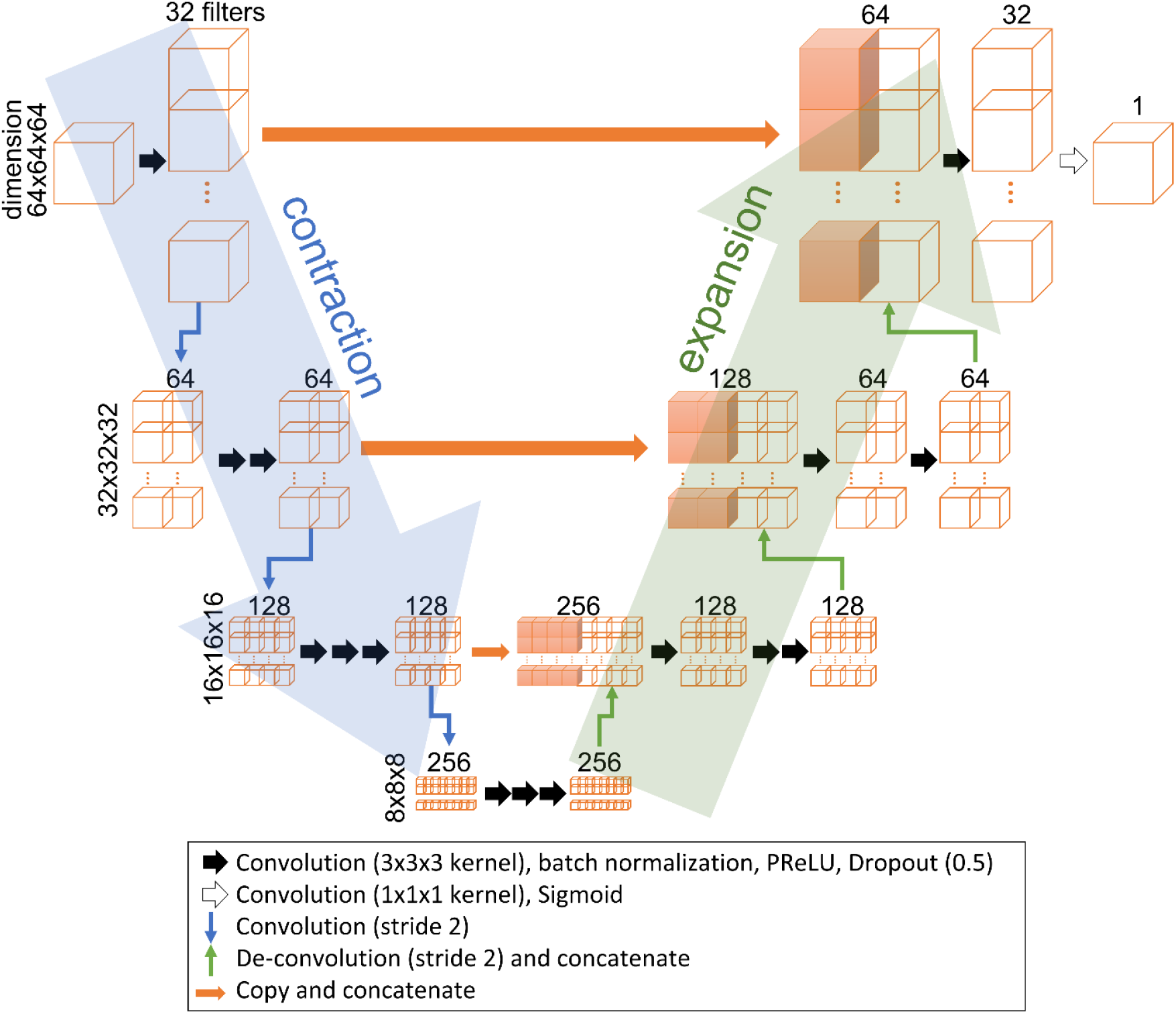
Schematic representation of the model network architecture. The left half is the contraction part reducing each volume dimension from 64×64×64 down to 8×8×8 voxels, and the right half is the expansion part enlarging the dimension up to the original volume, 64×64×64 voxels. The number of filters increases at the contraction part and decreases at the expansion part. The black arrows represent a set of 3-D convolution with 3×3×3 kernel size preserving the input dimension, batch normalization, PReLU activation function, and dropout operations. The white arrow represents a set of 3-D convolution with 1×1×1 kernel size and Sigmoid activation function operations. The downward blue arrows represent 3-D convolution with stride 2 to half the image size. The upward green arrows represent 3-D convolution with stride 2 to double the image size. The orange arrows represent copy operation. The results of green and orange operations are concatenated.

Our V-Net model was implemented in Keras 2.4.3 (Chollet and others, 2015) with tensorflow-gpu 2.2.0 (Martín Abadi et al., 2015) and CUDA 10.1 (NVIDIA, 2019) backend, modified from a V-Net implementation in GitHub (https://github.com/MiguelMonteiro/VNet-Tensorflow). The detailed parameters were: number of levels = 3, number of convolutions = (1, 2, 3), number of channels = (32, 64, 128), number of bottom convolutions = 3, activation function before the last layer = Parametric Rectified Linear Unit (PReLU), (He et al., 2015), dropout rate = 0.5, number of epochs = 50, optimizer = Adam (Kingma and Ba, 2017), initial learning rate = 0.001, loss function = dice similarity coefficient (DSC), (Dice, 1945). We trained the model on a NVIDIA^®^ TeslaTM T4 16GB system.

### 2.4. Prediction and evaluation

To apply the trained model to other datasets, they were first AC-PC-aligned by rigid-body registration to the MNI152 template, resampled to 0.7 mm isotropic resolution, and cropped to 64×64×64 voxels centered at the average Hb location of the training data. Then, the trained model predicted the Hb segmentation. We then applied automated postprocessing on predicted segmentation to remove incorrect segmentation apart from Hb by removing disconnected regions from the segmentation, keeping two large regions.

The evaluation of the data with T1w/T2w myelin content-based Hb segmentation (test data of HCP-YA S1200 and CamCAN data) included visual inspection, DSC, mean distance (MD, mean of surface distances from one segmentation’s surface voxels to the other segmentation), and Hausdorff distance (HD), (Aspert et al., 2002), while only visual inspection was done for the Mount Sinai 7T datasets without myelin content-based Hb segmentation.

### 2.5. MNI space transformation

To compare average Hb segmentation between myelin content-based segmentation and deep learning-based segmentation, we transformed the segmentation of the test data to the MNI152 space using nonlinear transformations from the HCP pipeline.

### 2.6. Data and code availability

HCP-YA S1200 data are available at https://www.humanconnectome.org/study/hcp-young-adult. CamCAN data are available at https://www.cam-can.org/index.php?content=dataset.

Mount Sinai 7T data would remain confidential and would not be shared due to data ownership.

The codes for training, testing, and prediction, as well as HCP-YA S1200 subject lists of training, validation, and test sets, are available in a GitHub repository (https://github.com/joowon-kim/hb_seg_vnet). The trained model used in this manuscript is available upon request due to its large size (>400 MB) to share via GitHub.

## 3. Results

### 3.1. Training of the V-Net model

The V-Net model training reached a steady state of loss function values (1 - DSC) less than 0.25 for both training and validation sets after about 30 epochs (Supplementary Fig 1). The training took about 30 min for each epoch, a total of 25 hours for 50 epochs. We chose the model with the lowest validation loss function value after the 30th epoch as our Hb segmentation model for the rest of the analysis (epoch 44, validation loss = 0.2048).

### 3.2. Test data and CamCAN data

The predicted Hb segmentation from the test data (n = 157) and CamCAN data (n = 441) showed good agreement with the label Hb segmentation (Table 1 and Fig. 3 and 4). The DSCs were higher than 0.6 except for 3 (1.9 %) of the test data and 10 (2.3 %) of CamCAN data. The MDs were less than 0.5 mm for all of the test data and 20 (4.8 %) of CamCAN data, while the HDs were less than 3 mm except for 4 (2.5 %) of the test data and 24 (5.4 %) of CamCAN data. The threshold values of these measures were from our previous study about the reproducibility of myelin content-based Hb segmentation (Kim et al., 2018). The similarity of the test data was better than CamCAN data as expected because both the training data and the test data were HCP-YA S1200 data which were from the same scanner and scan parameters.

**Table 1.** Mean ± standard deviation of label Hb volume and predicted Hb volume as well as their similarity comparison using dice similarity coefficient, mean distance, and Hausdorff distance from the test data and CamCAN data.

| | Test ($n = 157$) | CamCAN ($n = 441$) |
| --- | --- | --- |
| **Label volume ($\text{mm}^3$)** | $44.56 \pm 8.11$ | $49.64 \pm 13.01$ |
| **Prediction volume ($\text{mm}^3$)** | $45.53 \pm 3.94$ | $40.01 \pm 4.40$ |
| **Dice similarity coefficient** | $0.79 \pm 0.06$ | $0.71 \pm 0.05$ |
| **Mean distance (mm)** | $0.22 \pm 0.07$ | $0.34 \pm 0.09$ |
| **Hausdorff distance (mm)** | $1.63 \pm 0.50$ | $2.05 \pm 0.61$ |

**Figure 3.**
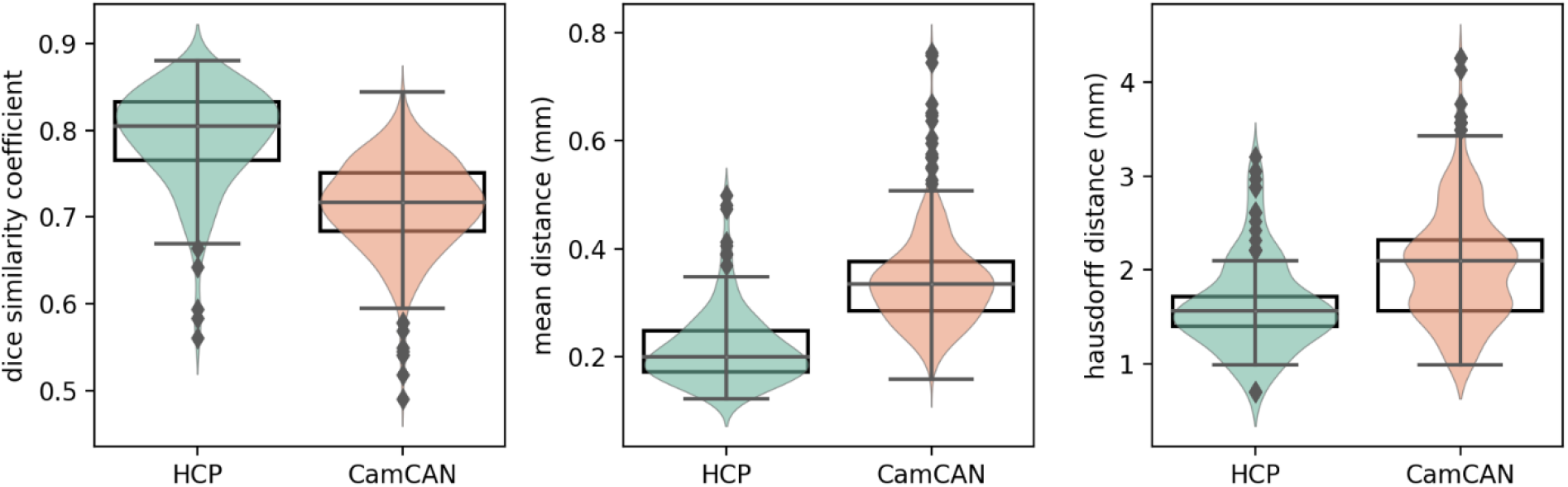
Box plots on top of violin plots of dice similarity coefficient, mean distance, and Hausdorff distance from the test data (n=157) and CamCAN data (n=441).

**Figure 4.**
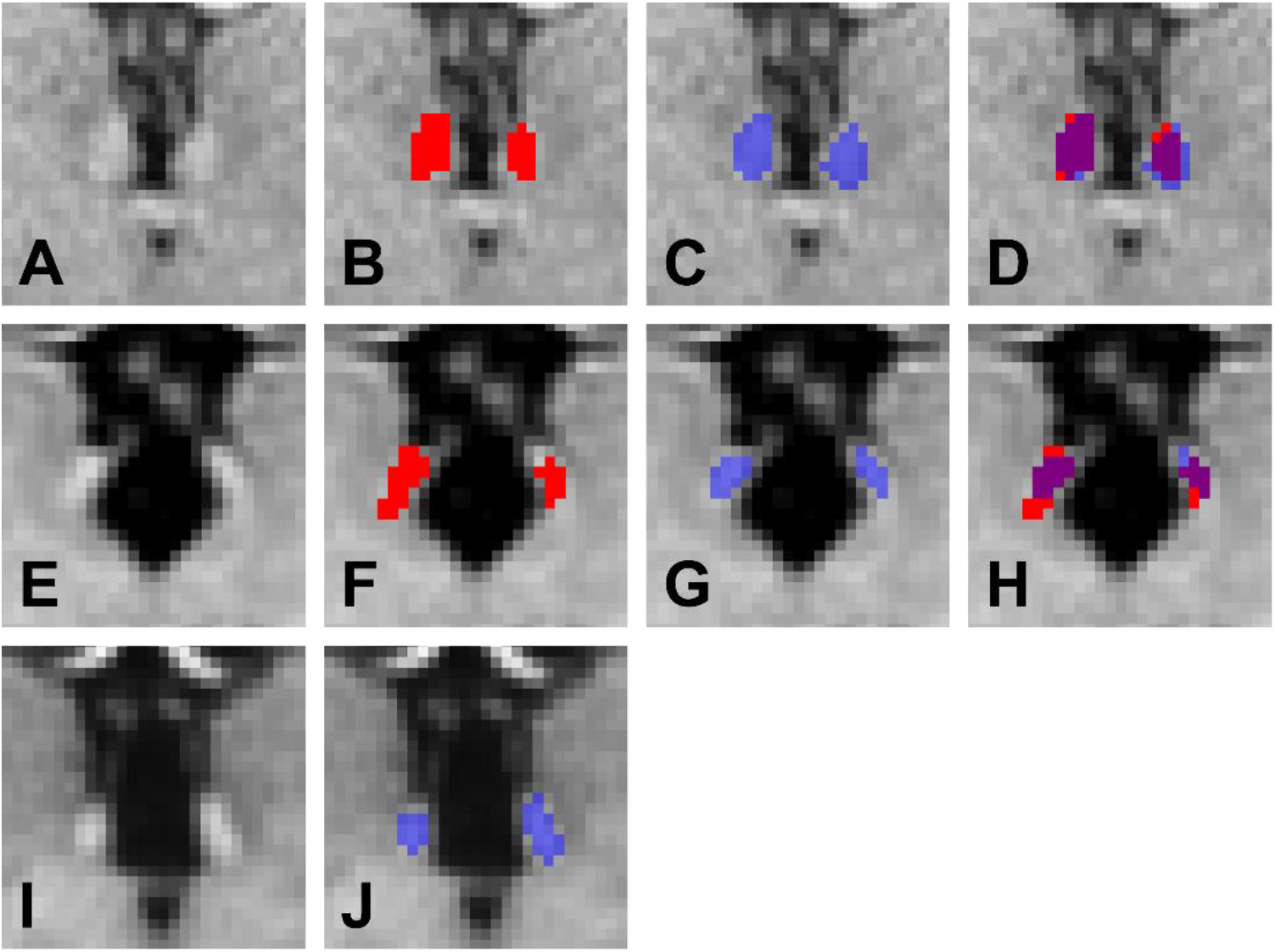
Representative coronal T1w images zoomed around Hb (A, E, I, and K) and Hb segmentation of label (B and F), prediction (C, G, J, and L), and the comparison between label and prediction (D and H) from a test subject (A-D, DSC=0.77, MD=0.24 mm, HD=1.21 mm), a CamCAN subject (E-H, DSC=0.74, MD=0.29 mm, HD=2.10 mm), and a Mount Sinai 7T subject (I-J). Red-colored regions are label Hb segmentation (B, D, F, and H), blue regions are prediction (C, D, G, H, and J), and purple regions are the intersection between label and prediction (D and H).

### 3.3. Other data and previously failed cases

The Hb was successfully segmented from Mount Sinai 7T data (n=5) (Fig. 4 I-J). For the subjects whose previous T1w/T2w myelin content-based segmentation failed due to insufficient contrast, the deep learning-based method successfully segmented the Hb (Supplementary Fig. 2).

### 3.4. MNI space transformation

The average Hb segmentation of the test data transformed to the MNI152 space is shown in Fig. X. Compared to the label segmentation (left, Fig. 5), the predicted segmentation (right, Fig. 5) was less over-estimated along the fasciculus retroflexus, a highly myelinated efferent fiber from the habenula (black arrow) and more overlapped across subjects.

**Figure 5.**
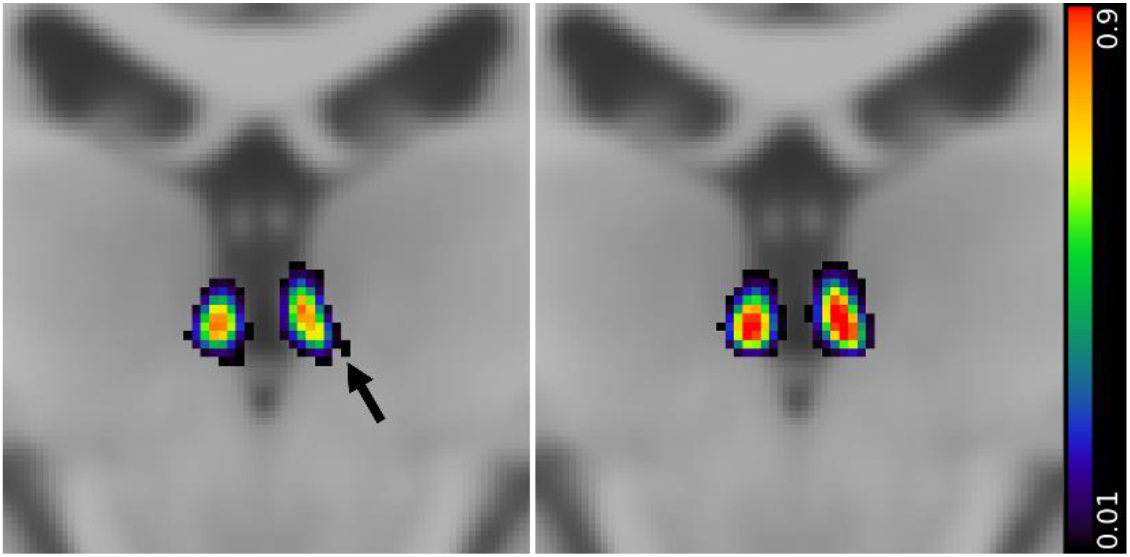
T1w MNI152 template images overlaid with average Hb label (left) and prediction (right) of the test data (n = 157) transformed to the MNI152 space. A black arrow points to the fasciculus retroflexus.

## 4. Discussion

We have demonstrated a deep learning-based Hb segmentation from T1w images that was automated, objective, fast, robust, and reliable. A large amount of readily available labels were from our myelin content-based segmentation that used all T1w, T2w, and T1w/T2w ratio images in our previous studies. On the other hand, the training was on only T1w images, leading to prediction available on T1w images alone. Without requiring T2w or T1w/T2w ratio images, the deep learning-based segmentation showed comparable performance - high DSC and short MD/HD - to the myelin content-based segmentation. No manual work is necessary during deep learning-based segmentation prediction except for visual inspection.

Unlike other deep learning-based segmentation, our approach used different inputs when training (T1w images alone) from those when creating labels (T1w, T2w, and T1w/T2w images). Due to this difference, we cannot expect identical or near-identical predictions to labels even in training sets. Nevertheless, our DSC, MD, and HD are comparable to other deep learning-based segmentation methods for spinal cord gray matter (Perone et al., 2018) or hippocampus (Carmo et al., 2021), which is more than expected.

We highly recommend a visual inspection of each subject’s segmentation to find obvious erroneous segmentation, although there were no such cases in our data. The postprocessing to remove off-Hb-region was necessary because the predicted Hb regions sometimes (less than 1 %) contain voxels far off the Hb region. After the postprocessing, visual inspection became straightforward to check 5 to 10 slices around the Hb region.

In addition to the high similarity in the test data, this new approach also showed robustness. It successfully segmented the Hb of the subjects that myelin content-based segmentation failed to segment Hb. We expect that this is not uncommon because myelin content-based segmentation requires high-quality, well-aligned T1w and T2w images; if either image quality is compromised by subject motion or the alignment is imperfect, the T1w/T2w image quality decreases, leading to unsatisfactory segmentation using the myelin content-based method.

When segmenting other datasets, we resampled input images to the same resolution as the training data. This resampling ensures similar sizes of inputs but introduces blurry effects. Using native resolution might be viable if we train various resolution data, but it is not straightforward to tell which way is preferred in terms of segmentation performance.

The training data were of young adults scanned with the same scanner and imaging parameters, leading to high performance on similar data types but lower performance on others. Adding another training data with a wide age range, multiple scanners, and different imaging parameters may increase the robustness of the segmentation.

The training labels were based on myelin content that is heterogeneous in Hb (Díaz et al., 2011). Therefore, the myelin content-based segmentation sometimes underestimates Hb near a less myelinated area but overestimates along highly myelinated fibers connected to the Hb (e.g., fasciculus retroflexus or stria medullaris). We did not observe group-wise overestimation along fasciculus retroflexus in the deep learning-based segmentation. This is likely due to the lack of T2w or T1w/T2w images, which accentuates the myelin contrast. This could indicate that the deep learning-based segmentation relies less on intensities but more on shapes than myelin content-based segmentation.

## Acknowledgment

We are grateful to Dr. Sophia Frangou (currently at University of British Columbia) at Icahn School of Medicine at Mount Sinai for sharing the Mount Sinai 7T data.

Data for deep learning were provided by the Human Connectome Project, WU-Minn Consortium (Principal Investigators: David Van Essen and Kamil Ugurbil; 1U54MH091657) funded by the 16 NIH Institutes and Centers that support the NIH Blueprint for Neuroscience Research; and by the McDonnell Center for Systems Neuroscience at Washington University.

Data collection and sharing for 1 mm isotropic resolution data was provided by the Cambridge Centre for Ageing and Neuroscience (CamCAN). CamCAN funding was provided by the UK Biotechnology and Biological Sciences Research Council (grant number BB/H008217/1), together with support from the UK Medical Research Council and University of Cambridge, UK.

