## Supplementary for "Automated Human Habenula Segmentation from T1-weighted Magnetic Resonance Images using V-Net"

**Supplementary Materials**

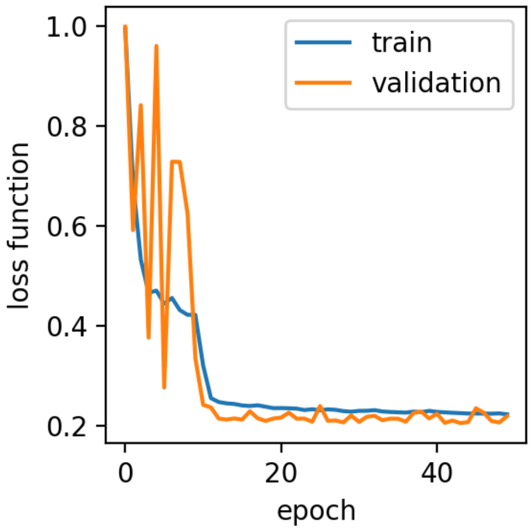


Supplementary Figure 1. Training and validation loss function (1 - dice similarity coefficient).

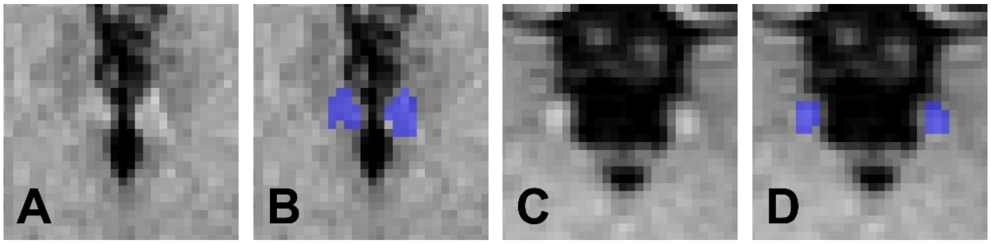


Supplementary Figure 2. Representative coronal T1-weighted images zoomed around Hb (A and C) and predicted Hb segmentation (B and D) from previously failed using T1-weighted/T2-weighted myelin content-based segmentation method HCP-YA S1200 subject (A-B) and CamCAN subject (C-D).
